# Real Time Face Recognition with limited training data: Feature Transfer Learning integrating CNN and Sparse Approximation

**DOI:** 10.1101/2021.03.17.435457

**Authors:** Supriya Bajpai, Gargi Mishra

## Abstract

It is highly challenging to obtain high performance with limited and unconstrained data in real time face recognition applications. Sparse Approximation is a fast and computationally efficient method for the above application as it requires no training time as compared to deep learning methods. It eliminates the training time by assuming that the test image can be approximated by the sum of individual contributions of the training images from different classes and the class with maximum contribution is closest to the test image. The efficiency of the Sparse Approximation method can be further increased by providing high quality features as input for classification. Hence, we propose to integrate pre-trained CNN architecture to extract the highly discriminative features from the image dataset for Sparse classification. The proposed approach provides better performance even for one training image per class in complex environment as compared to the existing methods. Highlight of the present approach is the results obtained for LFW dataset with one and thirteen training images per class are 84.86% and 96.14% respectively, whereas the existing deep learning methods use a large amount of training data to achieve comparable results.

## 1. Introduction

The ability of humans to recognize different faces has attracted many researchers to build machine learning models for face recognition (FR). The FR research focuses on the process of providing machines the ability to identify and verify facial images. The face recognition models learn to map the face image to a feature vectors and measure the distance corresponding to the face similarity. Despite the considerable improvement in performance of FR algorithms, good classification accuracy with limited data is still a challenge. The efficiency of FR algorithms is severely affected due to non linear variations present in unconstrained environment such as change in pose, expression, illumination or any additional physical variations such as scarf, glasses and beard.

Sparse representation method for classification performs well on small training datasets but the performance deteriorates with increase in image complexity. The deep learning models provides high performance for constrained as well as unconstrained datasets but training of these models requires huge amount of data and time. So the challenge emerges with real life data having limited samples and highly nonlinear variations due to unconstrained environment. The challenge posed by the complexity of space and time can be tackled by using transfer learning approach. Transfer learning refers to fine-tuning of an existing model or feature extraction from the layers of pre-trained deep neural network. In transfer learning approach the image feature vectors are extracted using the deep pre-trained neural networks and these feature vectors are transfered to other networks for training and classification.

The deep learning models such as convolution neural network (CNN) have the capability to handle the nonlinear complex facial variations. In CNNs, the initial layers are observed to learn the important features and the later layers provide learning of higher level abstractions [1]. These higher level abstractions represent facial identities with outstanding stability. Therefore, the feature extracted from the pretrained deep learning models such as CNN shows remarkable performance [13]. VGGF, VGG16, VGG19 [2, 3], AlexNet, ResNet-50 [4] are few commonly used pre-trained CNN models for feature extraction.

In this paper we propose a novel approach for FR via transfer learning that combines CNN with linear sparse approximation (LSA) for facial recognition. We extract the features of the face datasets from the deep CNN architecture (Inception-ResNet-v1) [5], pre-trained on two different databases (VGGFace2 [6] and CASIA-Webface [7]) separately and classify using linear sparse approximation.To investigate the performance of the proposed method the experiments are carried out systematically and extensively on six different standard datasets. The obtained accuracy is better even for one training image per class in unconstrained and complex environment as compared to the existing methods.

## 2. Literature Survey

Over the last few years the face recognition accuracy has drastically improved. Different FR methods have been evolved over time ranging from various statistical techniques [8, 9, 10] to deep learning methods [11, 12, 13]. The sparse approximation based methods are outperforming existing techniques constrained to limited training data in terms of classification accuracy and easier implementation [14, 15, 16]. In the last decade, many sparse approximation based methods have evolved that performs very well [17, 18, 19]. In these methods, the test vectors are approximated to linear sparse combination of training vectors and final matching contribution is calculated for further classification of test vector. The Kernel based sparse representation algorithms [20, 21], obtained by using the transformation of input space into high dimensional feature space, performs better than the conventional sparse approximation based methods. Lu et al. [22] proposed weighted sparse representation technique, which is based on combining the local information into sparse based approximation in a unique manner. Weighted group sparse representation technique [23], proposed by Xin et al. [23] combines the local information with group sparse based approximation for integration of the label information. In extended interval type-II and kernel based sparse representation method (KBSRM) [20], extended interval type-II fuzzy membership function is combined with Kernel sparse based approximation for FR. It extracts information that is hidden due to non-linear variation and pixel value overlapping. Hence, sparse based approximation provides very good results for small datasets but the performance is not satisfactory with unconstrained complex images.

The accuracy of recognition systems have been observed to depend heavily on the image feature extraction technique [24, 25]. Various feature extraction techniques are available in literature. Principal component analysis (PCA) [26], independent component analysis (ICA) [27] and other low-dimensional representation based techniques follow certain distribution assumptions and are the popular image feature extraction techniques. These methods fail to address the facial changes in uncontrolled environment. Many researchers attempted to address this problem using local feature extraction methods such as Gabor [9, 28], Local Binary Pattern (LBP) [25] and their variants - [29]. These methods provide robust performance due to the invariance property of local filtering but these handcrafted features display lack of distinctiveness and compactness. Therefore, learning based feature descriptors attracted researchers, in which learning of local filters is emphasized for improved distinctiveness and codebooks are learned for compactness. However, these representations are still unable to handle nonlinear complex facial variations.

The ability of deep learning methods to easily learn the rich and compact feature vectors from very large data-sets makes them very lucrative for face recognition applications [30, 31, 32, 33, 34, 35]. Following this, the researchers have applied the transfer learning approach that makes it easier to use these already trained deep learning models for FR [36, 37, 22]. Developing a CNN network from scratch requires massive amount of time and data. To avoid this, transfer learning approach is gaining popularity among researchers which saves a lot of time and resources by the use of pre-trained networks for feature extraction. In this approach the learned weights from the pre-trained network layers are used for feature extraction [38, 39, 12, 40, 41]. The above discussion shows that the conventional machine learning methods such as, sparse based representation method performs well with even with limited data but only on constrained data whereas the deep learning models perform fantastically well even on unconstrained data but the method requires large amount of training data. Therefore, there is a need to develop a model that combines both the deep learning models with the traditional models so that the model performs well on unconstrained data even with limited training samples.

## 3. Methodology

The basic framework of proposed methodology is given in Figure 1. In this framework, FR is carried out using transfer learning approach via two different modules: feature extraction using deep CNN and classification using linear sparse approximation. In the first module feature vectors of all images (training and testing) for a given dataset are extracted from the deep layer of the CNN. The second module receives training and testing images in the form of feature vectors and performs classification using linear sparse approximation. The two modules of propose no training time as observed in deep learning methods.d method are discussed in next two subsections.

**Figure 1:**
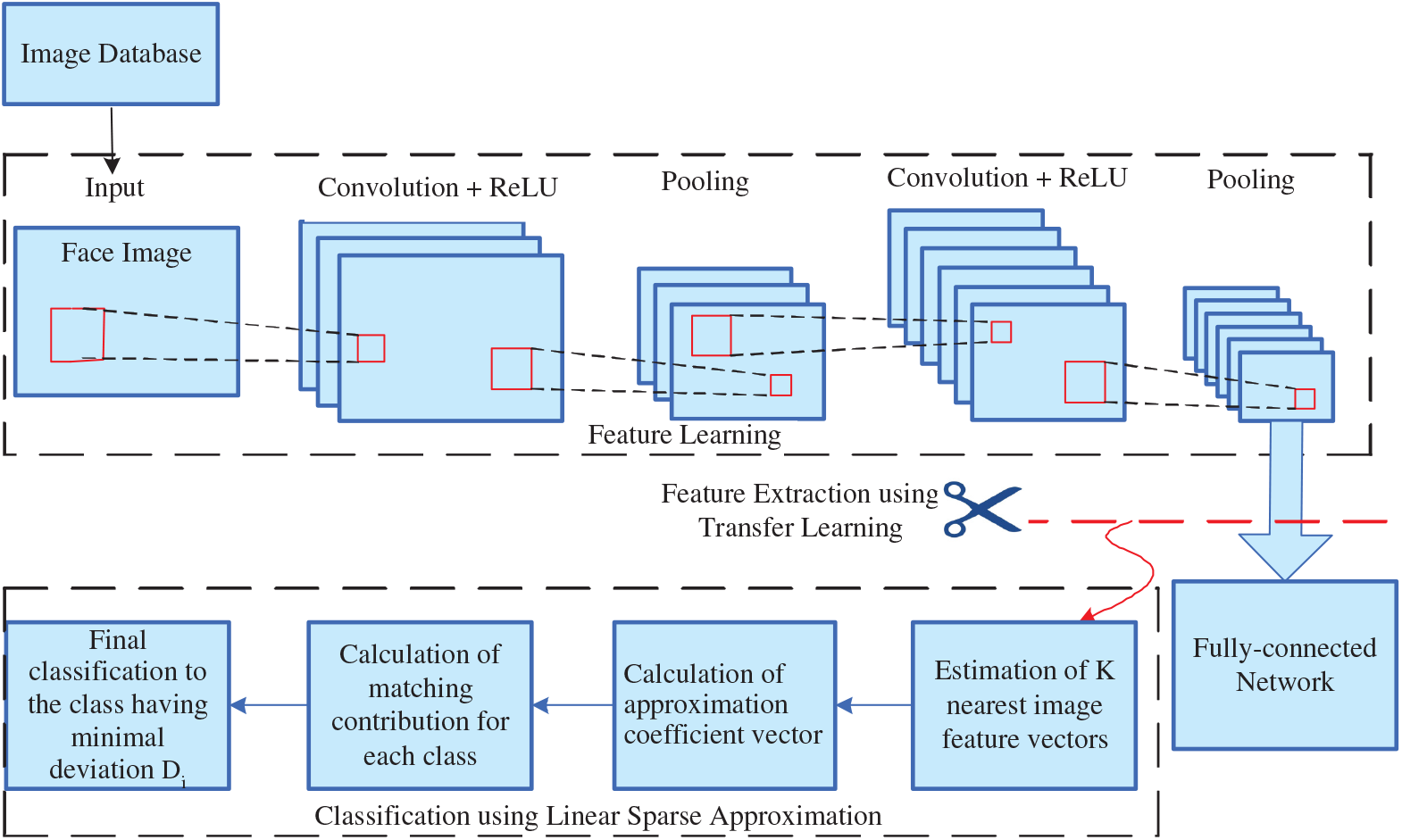
Basic framework of proposed method.

### 3.1. Module 1: Feature Extraction method using CNN

CNN networks have the capability to automatically learn different complex features from the images for different problems. The CNN architecture is composed of many layers of convolution, ReLU and max pooling, one or more fully connected layers and an output layer. The feature extraction is done by the convolution layers (convolution, ReLU and max pooling). During convolution the image is convolved with kernels/filters of same or different sizes. The mathematical expression for the convolution operation is given by

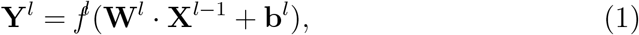

where, **W** is the kernel weights, **b** is the bias, *l* is the layer, **X** is the input feature map of *l* − 1^*th*^ layer and **Y** is the output feature map of *l*^*th*^ layer. The dimension of the obtained output feature maps after each layer is reduced by applying the pooling layer without changing the number of feature maps. The kernel weights **W** and bias **b** gets updated after each iteration during the training of the network via back propagation method.

In deep CNN each input image is represented as tensor **X** of size [H W C], where H, W, C is the height, width and number of color channels of image respectively. A pre-trained convolutional neural network (CNN) method forcan be represented as L number of functions in series *f*^1^, *f*^2^....*f*^*L*^ where L is the number of layers in the network. The output **Y**^*l*^ of layer *l* is given by (Eq. 1). The CNN learns layer weights and hence features through training the network and each layer learns different features. The deeper layers have the ability to learn complex features. The learned features are then used to classify the images. Image features can be computed from any layer of the pre-trained CNN by providing function *f*^*l*^, learned weights **W**^*l*^ and image tensor **X** such that, **Y**^*l*^ = *f*^*l*^(**X**^*l*−1^: **W**^*l*^).

We use a deep pre-trained CNN architecture (Inception-ResNet-v1) (Figure 2) [5] for feature extraction that has computational cost approximately equal to Inception-v3. Inception model architectures show high performance at low computational cost. Training of Inception network gets accelerated when combined with residual network. Combining Inception network with ResNet network also solves the problem of exploding/vanishing gradients that is a very common problem in deep network architectures.

**Figure 2:**
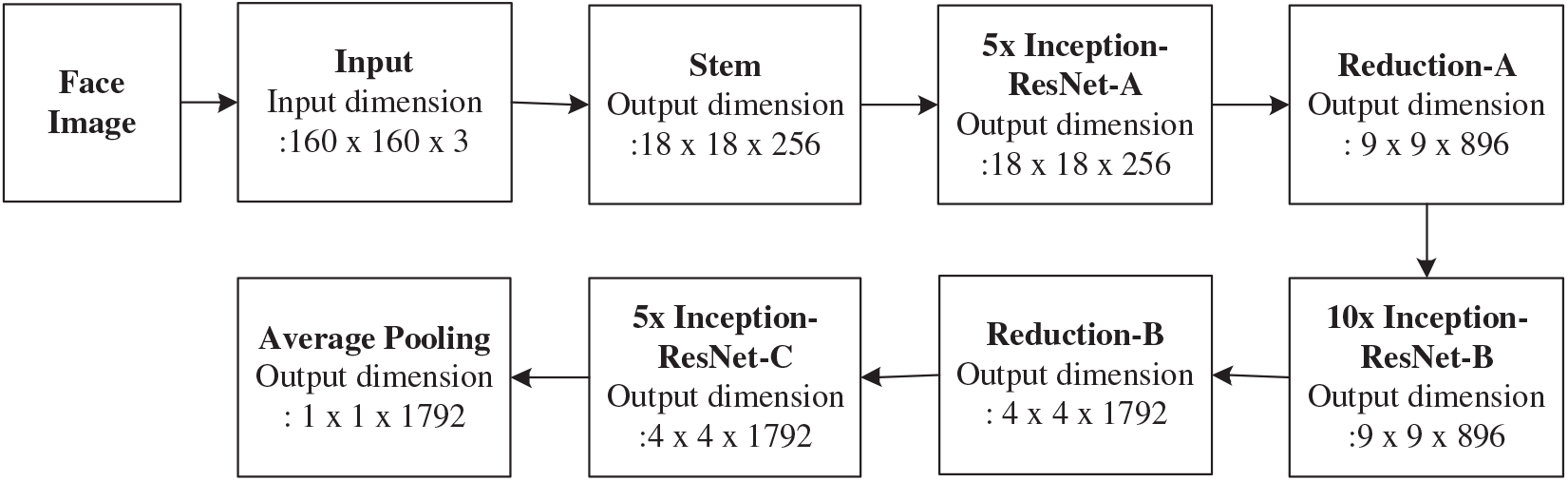
Schematic Representation of Inception-ResNet-v1 network.

In the present no training time as observed in deep learning methods.work, ResNet-Inception-v1 model [5] pre-trained with VGGFace2 [6] and Casia-Webfaces [7] database is used to extract the facial features. VGGFace2 is a large face database having a wide range of variations in pose, age, illumination, ethnicity and profession. The VGGFace2 dataset was proposed by Cao et al. consisting of 3.31 million face images of 9131 subjects, with an average of 362.6 images for each subject.

The feature vectors are extracted from the last layer (average pooling) of the network (Figure 1). The extracted feature vectors are then trained and classified using linear sparse approximation.

### 3.2. Module 2: Classification using Linear Sparse Approximation

Linear Sparse Approximation is a fast and computationally efficient classification method for face recognition as no previous training is required as compared to huge training requirements of deep learning methods. It eliminates the training time by assuming that the test image can be approximated by the sum of individual contributions of the training images from different classes and the class with maximum contribution is closest to the test image. Thus, to increase the efficiency of the classification highly discriminative features extracted from the above mentioned pre-trained CNN architecture is provided as input to the Sparse classifier.

This module receives all images of a given dataset in the form of feature vectors. Here, classification is implemented in three steps: in the first step, one nearest feature vector is identified from each class which is nearest to the test feature vector. Thus, the number of nearest feature vectors identified is equal to the number of classes i.e.′*K*′. In the second step, test feature vector is represented as linear sparse combination of identified ′*K*′ nearest feature vectors and coefficient values required for sparse approximation are calculated. In the third step, matching contribution for all nearest feature vectors is calculated. And finally the test feature vector is classified in to the class of nearest training feature vector having minimal deviation between its matching contribution and test feature vector.

#### Step 1 Determination of nearest feature vector

In this step, one nearest training feature vector is identified from each class using squared euclidean distance. Hence, total ′*K*′ nearest feature vectors are identified from complete training set. The elements of training set are represented by 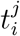 (i.e. *i*^*t*^*h* training feature vector from the *j*^*t*^*h* class, where *i* = 1, 2*, ..., T* and *j* = 1, 2*, ..., K*) and elements of testing set are represented by *q*_*r*_ (where *r* = 1, 2*, ...,* (*N − T*) × *K*). The *K* nearest training feature vectors of *q*_*r*_ are estimated using formula given in Equ. (2),

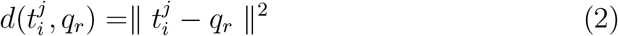

**Algorithm 1.**
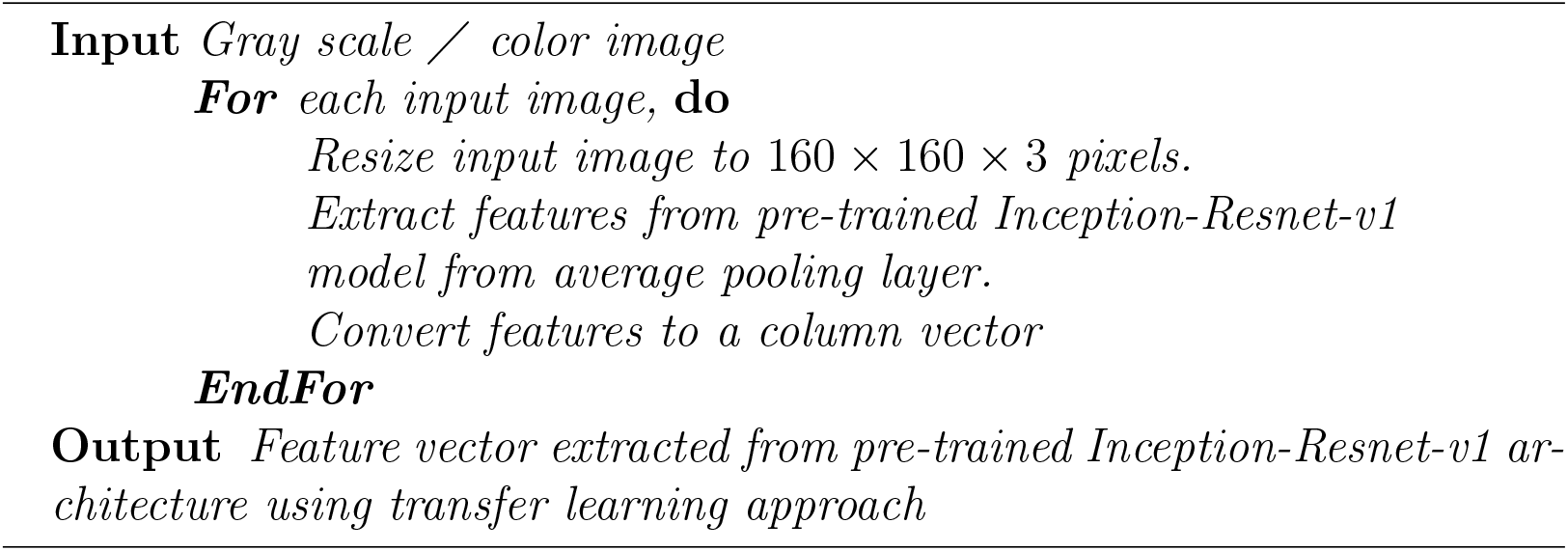
Feature vector extraction using pre-trained CNN model

For each class, distance between test feature vector and ′*T*′ training feature vectors is calculated which is denoted as *d*_1_*, d*_2_*, ..., d*_*T*_. Using these distances one nearest training feature vector having minimum distance is selected. Therefore, selecting one nearest feature vector from each class of training set makes a collection of ′*K*′ nearest feature vectors. Finally, all the ′*K*′ nearest training feature vectors *t*_1_*, t*_2_*, ..., t*_*K*_ are represented by a matrix *NTFV* = [*t*_1_*, t*_2_*, ..., t*_*K*_].

#### Step 2 Coefficient(α) calculation for linear sparse approximation

In this step, test image *q* is approximated as linear sparse combination of selected nearest training feature vectors. Here, it is assumed that the following equation is perfectly satisfied.

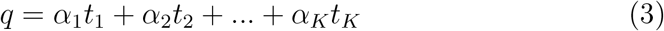

In Eq.(3), *t*_*m*_(*m* = 1, 2, ..., *K*) are *K* nearest training feature vectors and *α*_*n*_(*n* = 1, 2*, ..., K*) are the corresponding coefficients required for sparse approximation. In other words,

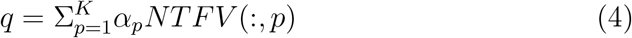

Eq.(3) can be rewritten in matrix form as

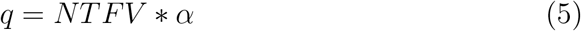

Where, *NTFV* = [*t*_1_*, t*_2_*, ..., t*_*K*_] and *α* = [*α*_1_*, α*_2_*, ..., α*_*K*_]^*T*^. Also the value of *α* is restricted to be the real number between −1 to +1 satisfying the condition *α*_1_ + *α*_2_ + ... + *α*_*K*_ = 1. Further, singularity test is performed on *NTFV*^*T*^*NTFV*. In case, it is found to be non singular, the Eq. (4) is solved using the formula given in Eq. (5).

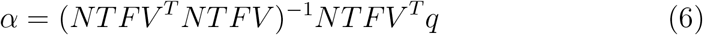

Otherwise, if *NTFV*^*T*^*NTFV* is nearly singular, *α* can be solved using Eq. (6).

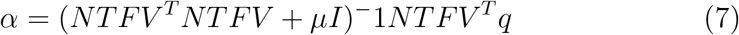

Here, *I* is the identity matrix and *μ* is a positive real number. Following the previous applications of sparse representation in face recognition, value of *μ* is set to 0.01. Therefore, coefficient *α* values are obtained using Eq. (5) and (6).

**Algorithm 2.**
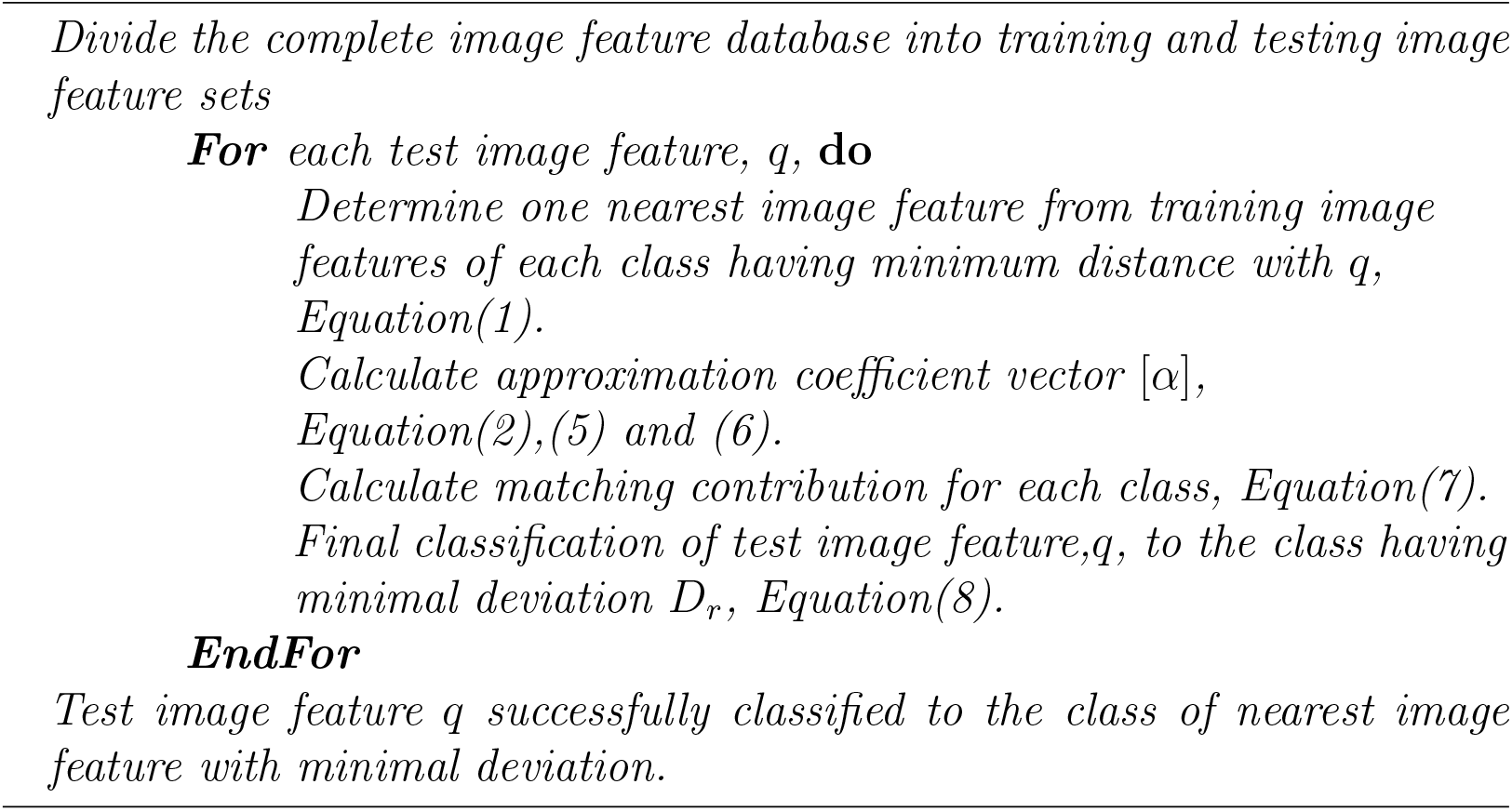
Classfication algorithm using linear sparse approximation

#### Step 3 Calculation of matching contribution and classification

It is clear from previous discussion that each nearest training feature vector present in *NT FV* is taken from a different class. For final classification, matching contribution of each class (or nearest training feature vector present in *NTFV*) is calculated. For *i*^*th*^ nearest training feature vector, matching contribution is calculated using Eq. (7) as,

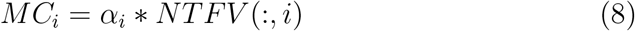

Matching contribution is obtained for each nearest training feature vector present in *NTFV*. The deviation between matching contribution of *r*^*t*^*h* nearest training feature vector and the test feature vector *q* is calculated using norm-2 distance as given in Eq. (8).

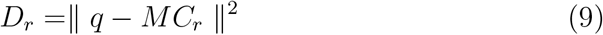

Here, it is clear that smaller value of *D*_*r*_ represents greater matching of *r*^*th*^ nearest training feature vector with test feature vector.

Hence, the test feature vector is classified into the class of nearest training feature vector having appropriate matching contribution and least deviation with test feature vector *q*. The complete algorithm of proposed method including sufficient programming details is given in Algorithm 1 and 2.

## 4. Experiments and results

In the present study we evaluate the performance of the proposed method with limited number of training images in unconstrained environment. by performing two different experiments. These experiments examine the variation in FR performance on various datasets and highlight the performance differences when CNN architecture (Inception-ResNet-v1) is pre-trained with VGGFace2 and CASIA-Webface. In the first experiment, the classification accuracy of the proposed model is evaluated on different datasets when image features vectors are extracted from Inception-ResNet-v1 pre-trained with VGGFace2 database. The second experiment evaluates the performance when Inception-ResNet-v1 is pre-trained with CASIA-Webface.

To perform the experiments, the extracted feature vectors are divided into mutually exclusive training and testing feature sets. One feature vector corresponds to one image. For each dataset, *T* feature vectors out of *N* feature vectors per class are selected as training features and remaining as (*N − T*) feature vectors per class are testing features, with a total number of possible training sets as 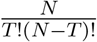. The details of total number of tests conducted for each dataset is given in Table 1. Here, it is to be noted that the test and training data being used in different experiments are mutually exclusive that avoids any possibility of overfitting. The results are expressed in terms of mean percentage classification accuracy (MPCA), maximum accuracy, minimum accuracy and standard deviation for all possible combination of training images per class (TIPC). To calculate the MPCA, if *q*_*1*_*, q*_2_*, ...q*_(*N* −*T*)_ are the members of test feature set and *C*_*L*_ = 1, 2*, ...K* is the set of class labels. Assume that, 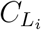 represents true class label of *q*_*i*_. For the proposed classifier *f*, 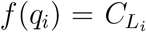 is the label prediction for test feature vector *q*_*i*_ where, 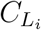 is a number from set *C*_*L*_. The mathematical expression for MPCA is given as:

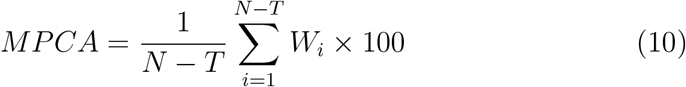

where,

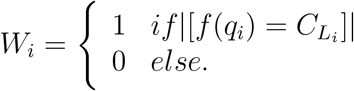

**Table 1:** Details of total tests conducted for each dataset.

| Dataset | Training images<br>per class (TIPC) | Number of<br>test sets | Number of<br>tests/ set | Total number<br>of tests conducted |
| --- | --- | --- | --- | --- |
| ORL | 1 | 10 | 360 | 3,600 |
|  | 2 | 45 | 320 | 14,400 |
|  | 4 | 210 | 240 | 50,400 |
|  | 6 | 210 | 160 | 33,600 |
|  | 9 | 10 | 40 | 400 |
| YALE | 1 | 11 | 15 | 1,650 |
|  | 2 | 55 | 135 | 7,425 |
|  | 4 | 330 | 105 | 34,650 |
|  | 6 | 462 | 75 | 34,650 |
|  | 10 | 11 | 15 | 165 |
| AR | 1 | 26 | 1,625 | 42,250 |
|  | 2 | 325 | 1,560 | 5,07,000 |
|  | 4 | 14,950 | 1,430 | 21,378,500 |
|  | 6 | 2,30,230 | 1,300 | 29,92,99,000 |
|  | 12 | 96,57,700 | 910 | 8,78,85,07,000 |
|  | 25 | 26 | 65 | 1,690 |
| GT | 1 | 15 | 700 | 10,500 |
|  | 2 | 105 | 650 | 68,250 |
|  | 4 | 1,365 | 550 | 7,50,750 |
|  | 6 | 5,005 | 450 | 22,52,250 |
|  | 12 | 455 | 150 | 68,250 |
|  | 14 | 15 | 50 | 750 |
| FEI | 1 | 14 | 650 | 9,100 |
|  | 2 | 91 | 600 | 54,600 |
|  | 4 | 1,001 | 500 | 5,00,500 |
|  | 6 | 3,003 | 400 | 12,01,200 |
|  | 12 | 91 | 100 | 9,100 |
|  | 13 | 14 | 50 | 700 |
| LFW | 1 | 14 | 650 | 9,100 |
|  | 2 | 91 | 600 | 54,600 |
|  | 4 | 1,001 | 500 | 5,00,500 |
|  | 6 | 3,003 | 400 | 12,01,200 |
|  | 12 | 91 | 100 | 9,100 |
|  | 13 | 14 | 50 | 700 |

### Description of datasets used

The details of all datasets used in the present work are given in this section. The brief specifications of datasets used are found in Table 2.

- ORL dataset: The ORL dataset [42] has a total of 400 face images with 10 images per class. The image resolution is 92 × 112 pixels. The dataset contains grey scale images with dark background, upright frontal position and a slight difference in facial expressions, lighting and pose.
- YALE dataset: The YALE dataset [43] has total 165 images with 15 subjects and 11 images of each subject. The images vary in expressions and with and without glasses. Image resolution is 220 × 175 pixels and are in gray scale.
- GT dataset: Georgia Tech dataset [44] consists of 750 images of 50 subjects and 15 colour images of each subject. Each image resolution is 640 pixels by 480 pixels. The images are captured at Centre for Signal and Image Processing at Georgia Institute of Technology with fussy background. We used the cropped images with resolution 131 pixels by 176 pixels. The images are captured with upright frontal and tilted pose with varying illumination condition, facial expressions and scale.
- AR dataset: AR dataset [45] has total of 1690 colour images of 65 subjects with 26 images per subject. The images vary in gender, facial expressions, illumination and occlusion. The image resolution is 165 pixels by 120 pixels. The database also contains images with black glasses and face scarf.
- FEI dataset: The FEI dataset [46] used in the present paper contains 700 Brazilian faces of 50 subjects with 14 persons per subject. The images are in color with resolution 640 pixels by 480 pixels. The images vary in facial expressions and pose and have a homogeneous white background.
- LFW dataset: Labelled Faces in the Wild (LFW) [47] is a large image dataset collected from internet consisting of total 13000 images with 5749 subjects. The number of images per subject is variable. The images in the dataset has large variability in pose, expression, age, origin, background and resolution. We randomly selected 700 images from 50 subjects with 14 images per subject from the original dataset and resized all the images with resolution 250 pixels by 250 pixels.

**Table 2:** Detailed Description of Standard Datasets Used in Our Experiments

| Database Details | ORL | YALE | AR | Georgia Tech | FEI | LFW |
| --- | --- | --- | --- | --- | --- | --- |
| No. of Classes | 40 | 15 | 65 | 50 | 50 | 50 |
| No. of Images per Class | 10 | 11 | 26 | 15 | 14 | 14 |
| Image Size | $92 \times 112$ | $220 \times 175$ | $165 \times 120$<br>$\times 3$ | $131 \times 176$<br>$\times 3$ | $640 \times 480$<br>$\times 3$ | $250 \times 250$<br>$\times 3$ |
| Total Instances | 400 | 165 | 1690 | 750 | 700 | 700 |

### 4.1. Experiment 1: CNN architecture (Inception-ResNet-v1) pre-trained with VGGFace2

The comparison of classification accuracy in terms of MPCA for the model pre-trained with VGGFace2 for all datasets is given in Table 3 (Figure 3). The MPCA for the proposed model with 1 and 2 TIPC is very high as compared to existing sparse based methods and CNN algorithms. It is observed that MPCA increases with increase in number of TIPC for all datasets. The highest MPCA 100%, is obtained for ORL, YALE and GT datasets with 9, 6 and 9 TIPC respectively. The high accuracy for these datasets is due to constrained face images with homogeneous background. The highest MPCA for AR and FEI datasets is 99.88% and 95.14% with 25 and 13 TIPC repectively. It is observed that MPCA for FEI dataset is better than AR dataset at low TIPC whereas the performance of AR dataset improves in comparison to FEI at large number of TIPC. This degradation in performance is due to presence of large occluded image portions present in AR dataset. The highest MPCA for LFW dataset is 96.14% with 13 TIPC which is higher than FEI. The reason for lower performance with FEI dataset is presence of relevant information in small portion of total image area. It is notable that the performance of proposed method with LFW dataset is higher than the performance of existing CNN architectures available in literature [4].

**Table 3:** Mean classification accuracy (%) of proposed method with VGGFace2 as pretrain dataset for feature extraction.

| Dataset | Number of Training Images Per Class (TIPC) |  |  |  |  |  |  |  |  |  |
| --- | --- | --- | --- | --- | --- | --- | --- | --- | --- | --- |
|  | 1 | 2 | 4 | 6 | 9 | 10 | 12 | 13 | 14 | 25 |
| ORL | 99.56 | 99.83 | 99.94 | 99.95 | 100 | NA | NA | NA | NA | NA |
| YALE | 99.82 | 99.92 | 99.99 | 100 | 100 | 100 | NA | NA | NA | NA |
| AR | 75.71 | 86.25 | 92.77 | 93.71 | – | – | 98.78 | – | – | 99.88 |
| GT | 99.34 | 99.79 | 99.96 | 99.98 | 100 | 100 | 100 | 100 | 100 | NA |
| FEI | 82.67 | 90.44 | 93.15 | 93.96 | 94.57 | 94.68 | 94.95 | 95.14 | NA | NA |
| LFW | 84.86 | 90.75 | 93.71 | 94.66 | 95.37 | 95.41 | 95.99 | 96.14 | NA | NA |

**Figure 3:**
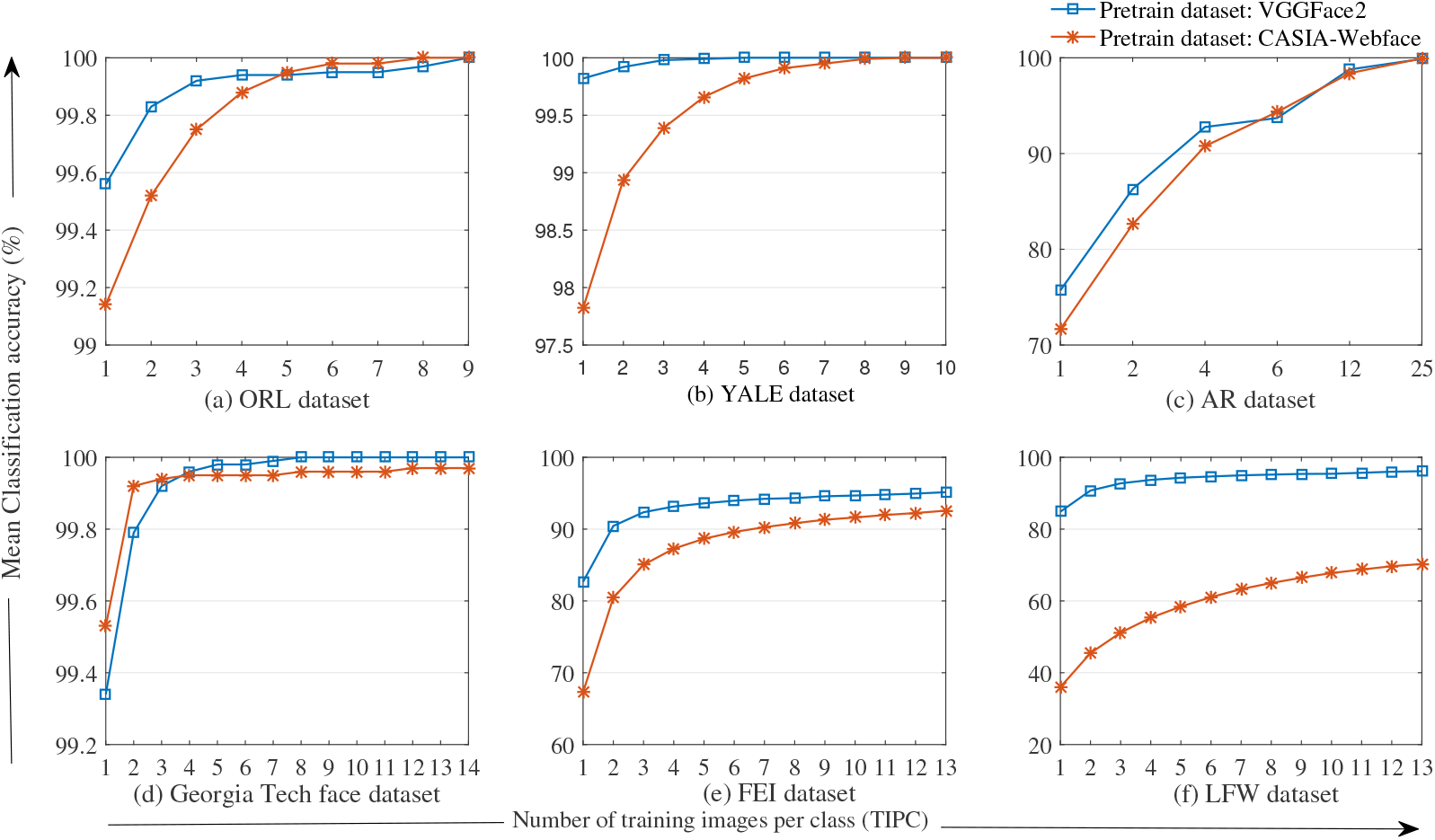
Variation in mean classification accuracy with number of TIPC for (a) ORL dataset (b) YALE dataset (c) AR dataset (d) Georgia Tech face dataset(e) FEI dataset (f) LFW dataset.

The statistical analysis of classification accuracy for all the tests conducted on datasets is performed in terms of maximum accuracy, minimum accuracy and standard deviation, shown in Figure 4 and Figure 5. From the plot, it is observed that for VGGFace2 the value of minimum accuracy is 100% at 9 TIPC for ORL, 5 TIPC onwards for YALE and 9 TIPC onwards for GT dataset. For AR, FEI and LFW datasets value of minimum accuracy is never achieved 100% for any number of TIPC. Similarly, the value of maximum accuracy is 100% for all possible number of TIPC of YALE and GT datasets and the value is 100% for 2 TIPC onwards for ORL, 12 TIPC onwards for AR and 13 TIPC onwards for FEI and LFW datasets.

**Figure 4:**
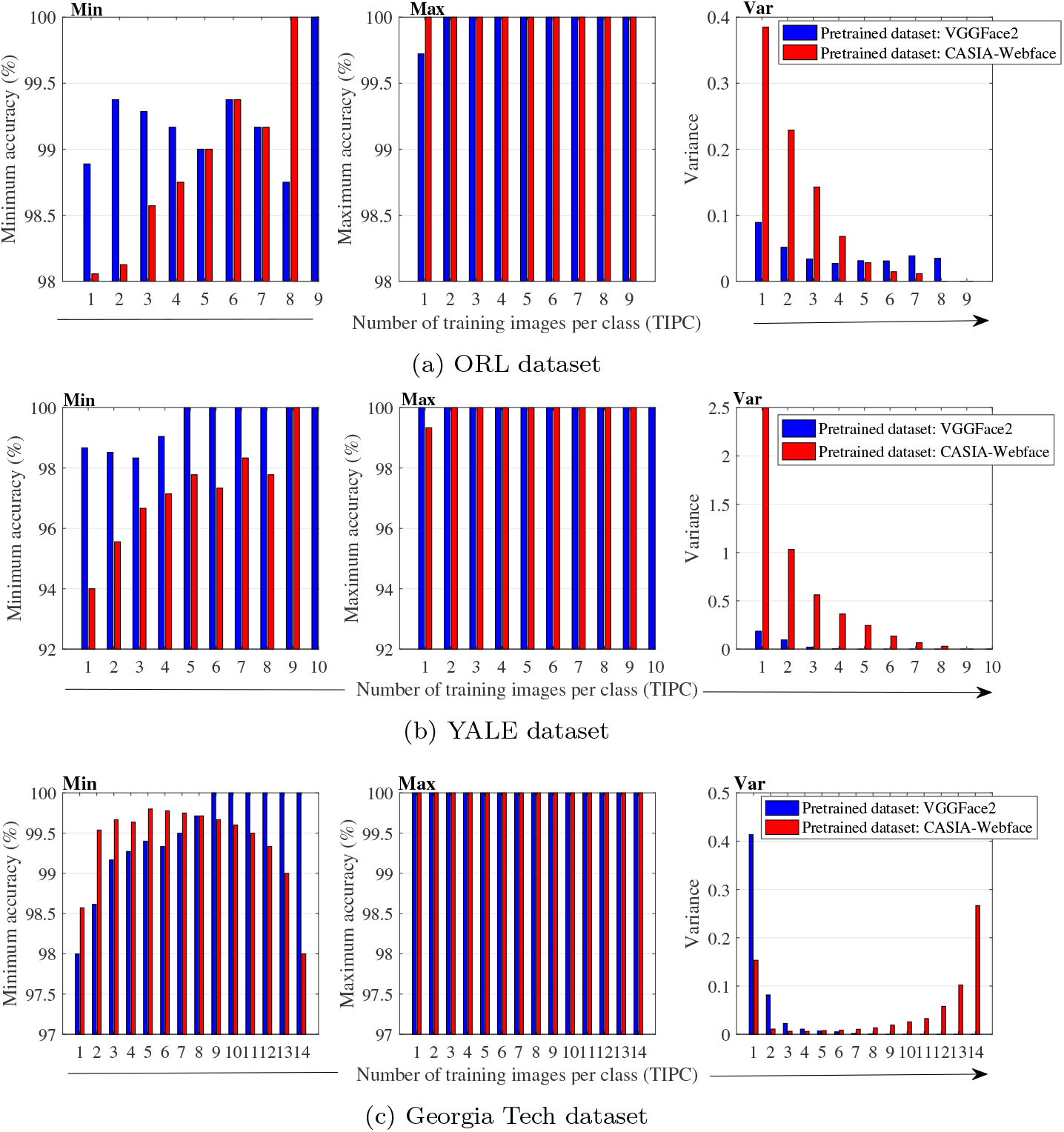
Statistical analysis of classification accuracy for (a) ORL dataset (b) YALE dataset (c) Georgia Tech dataset.

**Figure 5:**
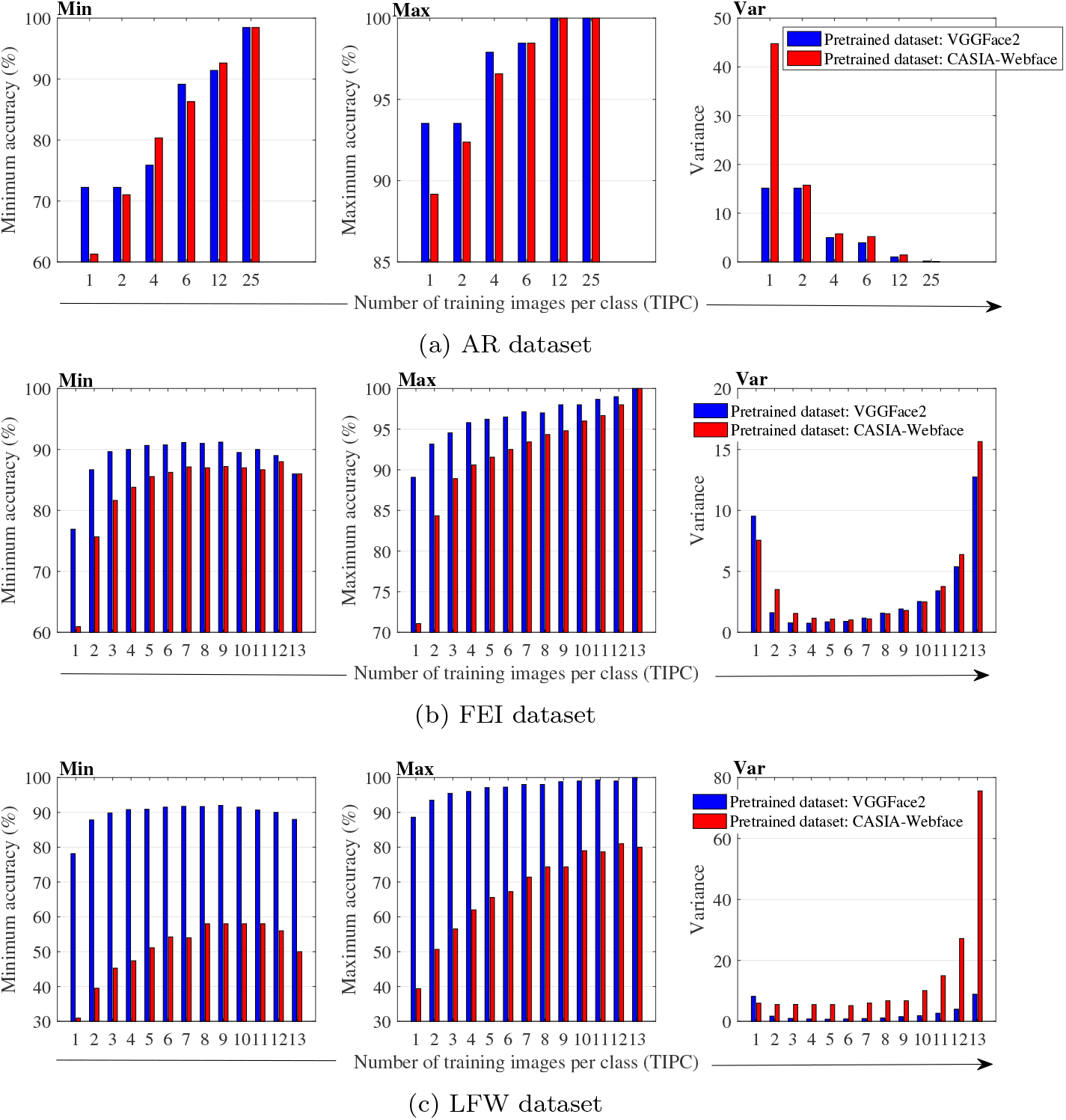
Statistical analysis of classification accuracy for (a) AR dataset (b) FEI dataset (c) LFW dataset.

### 4.2. Experiment 2: CNN architecture (Inception-ResNet-v1) pre-trained with CASIAWebface

The comparison of classification accuracy in terms of MPCA for the model pre-trained with CASIA-Webface for all datasets is given in Table 4 (Figure 3). The highest MPCA (100%) is obtained for ORL and YALE datasets at 9 and 10 TIPC respectively. The highest MPCA for AR, GT, FEI and LFW datasets is 99.94%, 99.97%, 92.57% and 79.29% with 25, 14, 13 and 13 TIPC respectively. It is observed that MPCA for LFW dataset is very low as compared to MPCA obtained in experiment 1, for all possible combinations of TIPC.

**Table 4:** Mean classification accuracy (%) of proposed method with CASIA-Webface as pre-train dataset for feature extraction.

| Dataset | Number of Training Images Per Class (TIPC) |  |  |  |  |  |  |  |  |  |
| --- | --- | --- | --- | --- | --- | --- | --- | --- | --- | --- |
|  | 1 | 2 | 4 | 6 | 9 | 10 | 12 | 13 | 14 | 25 |
| ORL | 99.14 | 99.52 | 99.88 | 99.98 | 100 | NA | NA | NA | NA | NA |
| YALE | 97.82 | 98.94 | 99.66 | 99.91 | 100 | 100 | NA | NA | NA | NA |
| AR | 71.65 | 82.6 | 90.77 | 94.35 | – | – | 98.39 | – | – | 99.94 |
| GT | 99.53 | 99.92 | 99.96 | 99.96 | 99.92 | 99.92 | 99.97 | 99.97 | 99.97 | NA |
| FEI | 67.29 | 80.44 | 87.28 | 89.60 | 91.31 | 91.61 | 92.24 | 92.57 | NA | NA |
| LFW | 36.01 | 45.47 | 55.32 | 61.04 | 66.46 | 67.77 | 69.67 | 70.29 | NA | NA |

The statistical analysis of classification accuracy for all the tests conducted on datasets is shown in Figure 4 and Figure 5. Plot shows that for CASIAWebface the value of minimum accuracy is 100% at 8 TIPC onwards for ORL and 9 TIPC onwards for YALE dataset. Also, for AR, GT, FEI and LFW datasets value of minimum accuracy is never achieved to be 100% for any number of TIPC. Similarly, the value of maximum accuracy is 100% for all possible number of TIPC of ORL and GT datasets and the value is 100% for 2 TIPC onwards for YALE, 12 TIPC onwards for AR and 13 TIPC onwards for FEI datasets. The value of maximum accuracy could not be achieved 100% for LFW dataset.

From the above experiments, it is observed that the overall performance in terms of MPCA for all datasets is better for model pre-trained with VGGFace2 database. Furthermore, in case of GT dataset performance is better at limited number of TIPC for model trained with CASIA-Webface whereas it is better at higher TIPC for model trained with VGGFace2. Also, the results obtained by the proposed model outperforms the existing methods for face recognition application (Table 5). One major constraint in comparing our results with the other published results in area of FR is the variability of training and testing datasets. The training dataset used in existing works using deep learning CNN models is extensively large as compared to our experiment training dataset.

**Table 5:** Comparison of the proposed method (S. Bajpai et al. (2020)) accuracy with other face recognition models.

| References | Datasets | Models | TIPC | MPCA |
| --- | --- | --- | --- | --- |
| S. Bajpai et al. (2020) | ORL | InceptionResnet-v1<br>+ LSA | **6** | 99.95% |
| S. Y. Wang et al. (2020) [38] | ORL | SAMPSR | 5 | 94.14% |
| S. Almadby et al. (2019) [4] | ORL | ResNet-50 + SVM | 8 | 100% |
| S. Guo et al. (2016) [38] | ORL | CNN+SVM | 7 | 97.5% |
| S. Bajpai et al. (2020) | YALE | InceptionResnet-v1<br>+ LSA | **6** | 100% |
| S. Zhu et al. (2017) [48] | YaleB-Extended | histogram based<br>feature representation |  | 99.37% |
| S. Bajpai et al. (2020) | Georgia Tech | InceptionResnet-v1<br>+ LSA | **12** | 100% |
| S. Bajpai et al. (2020) | AR | InceptionResnet-v1<br>+ LSA | **6** | 94.35% |
| S. Y. Wang et al. (2020) [38] | AR | SAMPSR | 8 | 72.49% |
| S. Bajpai et al. (2020) | AR | InceptionResnet-v1<br>+ LSA | **4** | 92.77% |
| N. Zhu et al. (2014) [20] | AR | KSR method | 4 | 91.61% |
| S. Bajpai et al. (2020) | FEI | InceptionResnet-v1<br>+ LSA | **13** | 95.14% |
| S. Bajpai et al. (2020) | FEI | InceptionResnet-v1<br>+ LSA | **1** | 82.67% |
| J. Cai et al. (2015) [49] | FEI | Sparse representation | 1 | 61.31% |
| S. Almadby et al. (2019) [4] | FEI | Transfer learning (AlexNet) | 11 | 98.7% |
| S. Bajpai et al. (2020) | LFW | InceptionResnet-v1<br>+ LSA | **13** | 96.14% |
| S. Bajpai et al. (2020) | LFW | InceptionResnet-v1<br>+ LSA | **10** | 95.41% |
| S. Almadby et al. (2019) [4] | LFW | ResNet-50 + SVM | 11 | 94% |
| S. Almadby et al. (2019) [4] | LFW | Transfer learning<br>(AlexNet) | 11 | 95.63% |

## 5. Conclusion

The paper presents an effective transfer learning approach for face recognition application combining pre-trained InceptionResnet-v1 deep CNN architecture and linear sparse approximation. The proposed method implements FR by extracting image features using InceptionResnet-v1 architecture and classifying using linear sparse approximation. The use of pre-trained CNN architecture for feature extraction improves the overall performance drastically by learning the higher level abstractions which represents facial identities with outstanding stability. Moreover, the classification using sparse based approximation presents less time complexity, ease of implementation and good accuracy with limited training data even with one and two TIPC. To examine the the performance of the proposed model two experiments are conducted on six different standard datasets using CNN architecture (Inception-ResNet-v1) pre-trained with two different datasets VGGFace2 and CASIA-Webface.

The experiment shows that the method performs better even in unconstrained environment with 1 and 2 TIPC as compared to the existing methods. In addition to that, the overall performance of the proposed method is better for the model pre-trained with VGGFace2 database.

